# THAP11_F80L_ cobalamin disorder-associated mutation reveals normal and pathogenic THAP11 functions in gene expression and cell proliferation

**DOI:** 10.1101/813352

**Authors:** Harmonie Dehaene, Viviane Praz, Philippe Lhôte, Maykel Lopes, Winship Herr

## Abstract

Twelve human THAP proteins share the THAP domain, an evolutionary conserved zinc-finger DNA-binding domain. Studies of different THAP proteins have indicated roles in gene transcription, cell proliferation and development. We have analyzed this protein family, focusing on THAP7 and THAP11. We show that human THAP proteins possess differing homo- and heterodimer formation properties and interaction abilities with the transcriptional co-regulator HCF-1. HEK-293 cells lacking THAP7 were viable but proliferated more slowly. In contrast, HEK-293 cells were very sensitive to THAP11 alteration. Nevertheless, HEK-293 cells bearing a THAP11 mutation identified in a patient suffering from cobalamin disorder (THAP11_F80L_) were viable although proliferated more slowly. Cobalamin disorder is an inborn vitamin deficiency characterized by neurodevelopmental abnormalities, most often owing to biallelic mutations in the *MMACHC* gene, whose gene product MMACHC is a key enzyme in the cobalamin (vitamin B_12_) metabolic pathway. We show that THAP11_F80L_ selectively affected promoter binding by THAP11, having more deleterious effects on a subset of THAP11 targets, and resulting in altered patterns of gene expression. In particular, THAP11_F80L_ exhibited a strong effect on association with the *MMACHC* promoter and led to a decrease in *MMACHC* gene transcription, suggesting that the THAP11_F80L_ mutation is directly responsible for the observed cobalamin disorder.

## Introduction

The THAP family of gene paralogs encodes 12 proteins in human, named THAP0 to THAP11. These are defined by their N-terminal THAP (for Thanatos — referring to the Greek God of Death — Associated Proteins) domain, an atypical zinc-finger DNA-binding domain [1,2]. THAP domains display similar three-dimensional structures, while recognizing different DNA target sequences [3–7]. Studies of different THAP proteins have indicated roles in gene transcription, cell proliferation and development. Here, we focus on two members: THAP7 and THAP11.

THAP7 represses gene transcription, both by promoting histone deacetylation and by masking histone tails from histone acetyltransferase complexes [8,9]. It is also implicated in the control of cell proliferation by abrogating Histone Nuclear Factor P (HiNF-P)-mediated activation of histone H4 gene transcription [10].

THAP11 (known as Ronin in mice, [11]) also regulates mammalian cell proliferation, with reported examples for both activation [11,12] and repression [13,14], and binds numerous promoters of genes involved in cell growth, metabolism, and cell cycle [12–17]. It also has essential transcriptional roles in pluripotency [11,18,19], hematopoiesis [20], and early development of retina [21], heart [17], and brain [22]. Furthermore, THAP11 is important for mitochondrial function by regulating nuclear mitochondrial-related genes, including components of the electron transport chain [21]. Consistent with the frequent involvement of THAP proteins in human diseases (e.g., dystonia 6 (THAP1, [23–25]), heart disease (THAP5, [26]), or cancers [7,12–14,27–32]), a *THAP11* missense mutation is associated with cobalamin disorder, an inborn vitamin deficiency characterized by neurodevelopmental abnormalities [22].

Different human mutations can result in disorders of intracellular cobalamin (vitamin B_12_) metabolism, defining several complementation groups. The *cblC* group, accounting for 80% of cobalamin-disorder patients, is owing to biallelic mutations in the *MMACHC* gene, whose gene product MMACHC is a key enzyme in the cobalamin metabolic pathway [33]. Interestingly, THAP11, together with its co-regulator HCF-1 (for herpes simplex virus Host Cell Factor 1), regulates the transcription of the *MMACHC* gene [5,22]. Indeed, certain mutations in the X-linked *HCFC1* gene encoding HCF-1 are responsible for the so-called *cblX* complementation group and result in both decreased *MMACHC* mRNA and MMACHC protein levels [33]. A cobalamin-deficiency patient has been recently described with a clinically *cblX*- like disorder but without either any *MMACHC* or *HCFC-1* mutation. Instead, this patient had a homozygous missense mutation in the *THAP11* gene resulting in a phenylalanine to leucine change at position 80 (referred to here as THAP11_F80L_) at the C-terminus of the THAP domain [22,33] — the causal link between this mutation and the cobalamin disorder, however, remains to be established.

Here, we analyze THAP7 and THAP11 interactions and probe their roles in cell proliferation, including through the study of the THAP11 F80L mutation.

## Results

### Structure of the human THAP family of proteins

To initiate the study of human THAP proteins, we examined their structural and evolutionary relationships. Fig 1 shows an evolutionary tree of the 12 human THAP proteins sorted by THAP-domain sequence similarity shown in S1 Fig. Although sharing the structurally-similar THAP domain, the distinct THAP proteins display considerable variability with sizes extending from 213 to 903 amino acids. Two other features shared by most of them are a so-called ^D^/_E_HxY HCF-1-Binding Motif (HBM; [34,35]) sequence for HCF-1 interaction (Fig 1, orange and dashed-orange lines) and a coiled-coil domain involved in protein homo- and heterodimer formation [36]. It has been argued that all [4] or all but one (THAP10) [37] of the 12 THAP proteins contain a coiled-coil domain. Nevertheless, in our analysis with two independent prediction tools (COILS [38] and PairCoil2 [39]), we detected a coiled-coil domain in only nine of the 12 THAP proteins; we did not detect a coiled-coil domain in THAP0, 9 and 10 (Fig 1, blue boxes). These two motifs, HBM and coiled-coil, appear together in seven of the 12 THAP proteins with the HBM invariably within 1 to 13 amino acids N-terminal of the coiled-coil domain, suggesting a functional evolutionary relationship.

**Fig 1.**
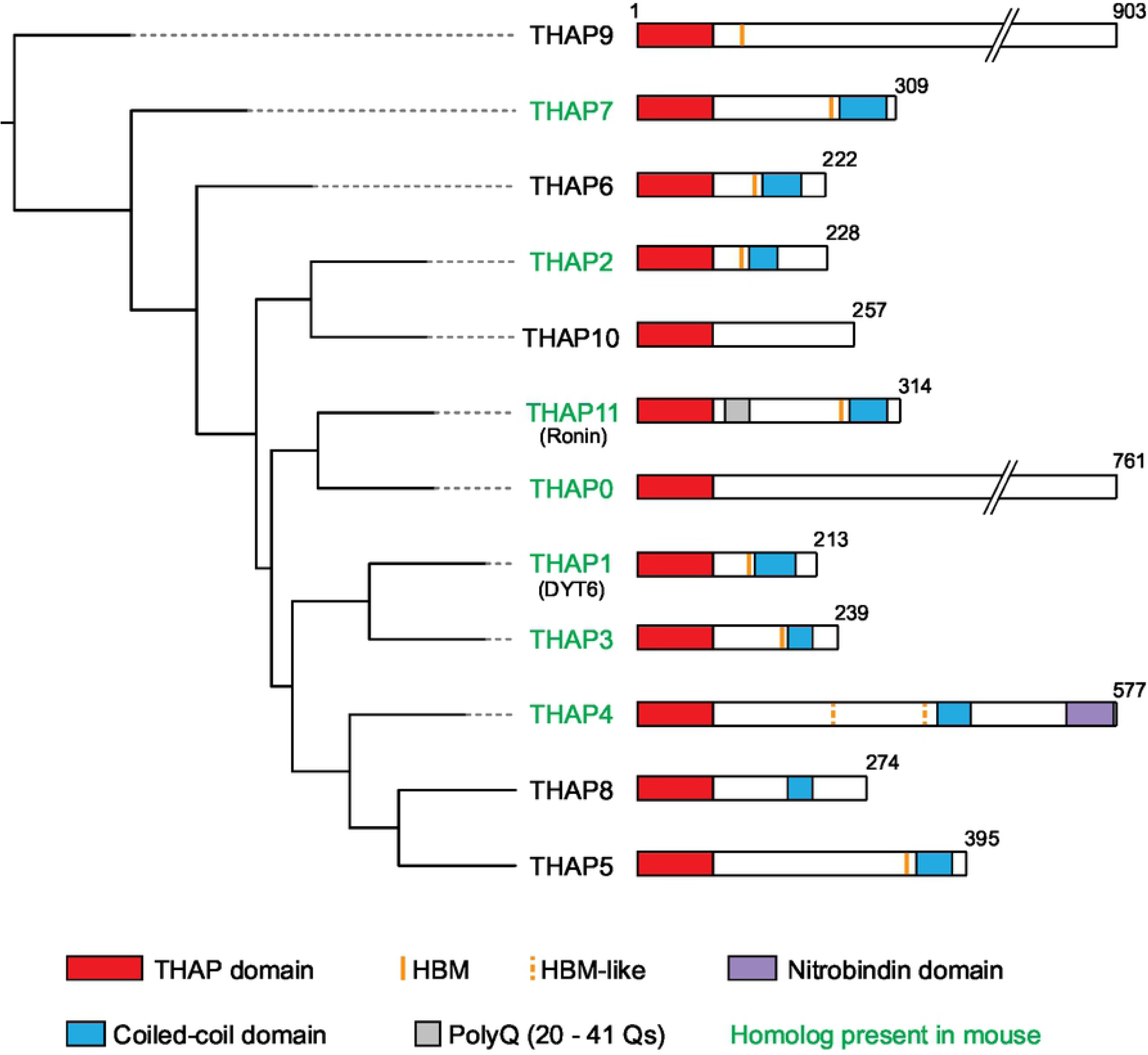
Evolutionary tree of the 12 human THAP proteins. THAP proteins are sorted by sequence similarity of their respective THAP domain. Sequence analyses were as described in S1 File. Sequence features are as indicated (see also text). Alternate names are shown in parenthesis and THAP proteins with homologs in mice are labelled in green.

In an extended study of THAP proteins [40], we analyzed the propensity for homodimer and heterodimer formation among THAP4, THAP5, THAP7, THAP8 and THAP11 proteins as well as their abilities to associate with HCF-1 via the HBM. In the context of this study, we decided to focus on two THAP proteins broadly expressed (see [40]) and having reported roles in transcriptional regulation [8,9,11]: THAP7 and THAP11.

### THAP7 and THAP11 form homodimers but not heterodimers

We first probe the formation of THAP7 and THAP11 homodimers and THAP7/THAP11 heterodimers. For this purpose, HEK-293 cells were co-transfected with Flag- and/or HA-epitope-tagged *THAP7* and *THAP11* expression constructs, and whole-cell lysates (wcl) subjected to anti-HA-epitope immunoprecipitation and analyzed by immunoblot. In Fig 2A, we test the ability of THAP7 to form homodimers. Interestingly, the ectopically synthesized THAP7 protein appears as a series of bands upon immunoblotting (lane 3, lower panel), probably owing to phosphorylation (S2A Fig). As evidenced by the recovery of Flag-tagged THAP7 protein after immunoprecipitation of co-synthesized HA-tagged THAP7 protein (compare lanes 8 and 4, lower panel), THAP7 forms homodimers effectively. Interestingly, the slower migrating form — and probably most heavily phosphorylated — is less effectively recovered than the faster migrating forms, suggesting that phosphorylation modulates THAP7 homodimer formation.

**Fig 2.**
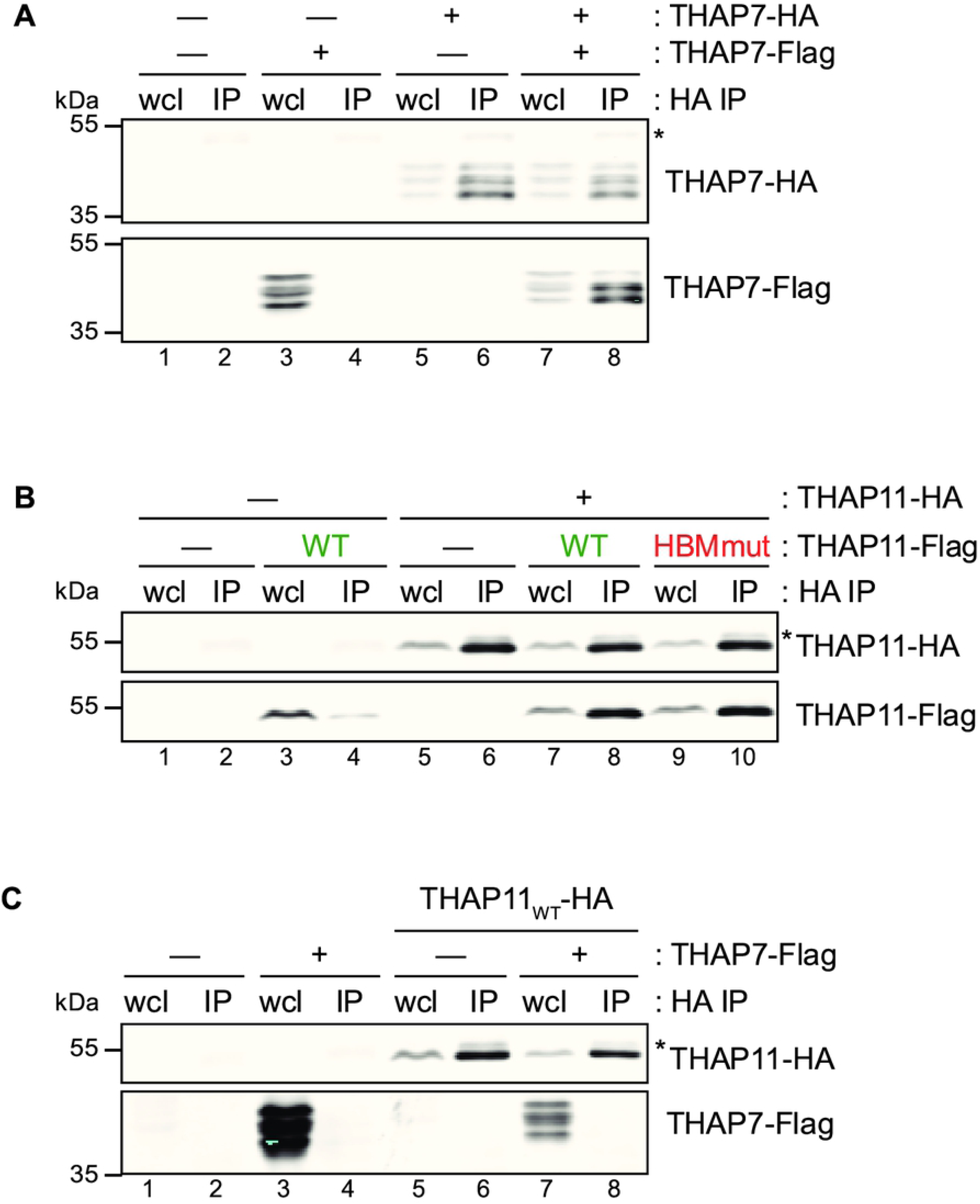
THAP7 and THAP11 homodimer but not heterodimer formation. HEK-293 cells were co-transfected with or without Flag- and HA-tagged THAP constructs (as indicated), and whole-cell lysates subjected to HA immunoprecipitation and analyzed by immunoblot with anti-HA (upper panels) and anti-Flag (lower panels) antibodies. (A) THAP7 homodimer formation. (B) THAP11 homodimer formation. HBM-positive THAP proteins are labeled in green whereas HBM-negative ones are in red. (C) THAP7 and THAP11 heterodimer formation. *, antibody heavy chain; wcl, whole-cell lysate; IP, immunoprecipitate.

As shown in Fig 2B, THAP11 protein also forms homodimers in this assay (compare lanes 4 and 8, lower panel), consistent with previous reports [11,41]. Although each forms homodimers, THAP7 and THAP11 do not form heterodimers as evidenced by the failed recovery of Flag-tagged THAP7 after immunoprecipitation of co-synthesized HA-tagged THAP11 (compare lanes 4 and 8 of Figs 2A and C). The segregated abilities of THAP7 and THAP11 to form homodimers but not heterodimers suggest that, although involved in similar transcriptional and cell proliferation processes, they do not directly influence each other’s transcriptional networks via dimer formation.

### THAP7 and THAP11 HBM sequences enable them to associate with HCF-1

Using an analogous epitope-tagged protein association assay as in Fig 2, we probed THAP7 and THAP11 association with HCF-1 as shown in Fig 3. HCF-1 is proteolytically processed into associated N- (HCF-1_N_) and C- (HCF-1_C_) terminal subunits [42,43]. As HCF-1 associates with HBM-containing proteins via its N-terminal Kelch domain [44], we tested THAP7 and THAP11 association with HCF-1_N_. As shown in Fig 3, both wild-type THAP7 and THAP11 associate with wild-type HCF-1_N_ as indicated by the recovery of HA-tagged HCF-1_N_ after immunoprecipitation of co-synthesized Flag-tagged THAP7 (Fig 3A) and THAP11 (Fig 3B) (compare lanes 4 with lanes 2; recovery of endogenous THAP11 in such an experiment is shown in S2B Fig).

**Fig 3.**
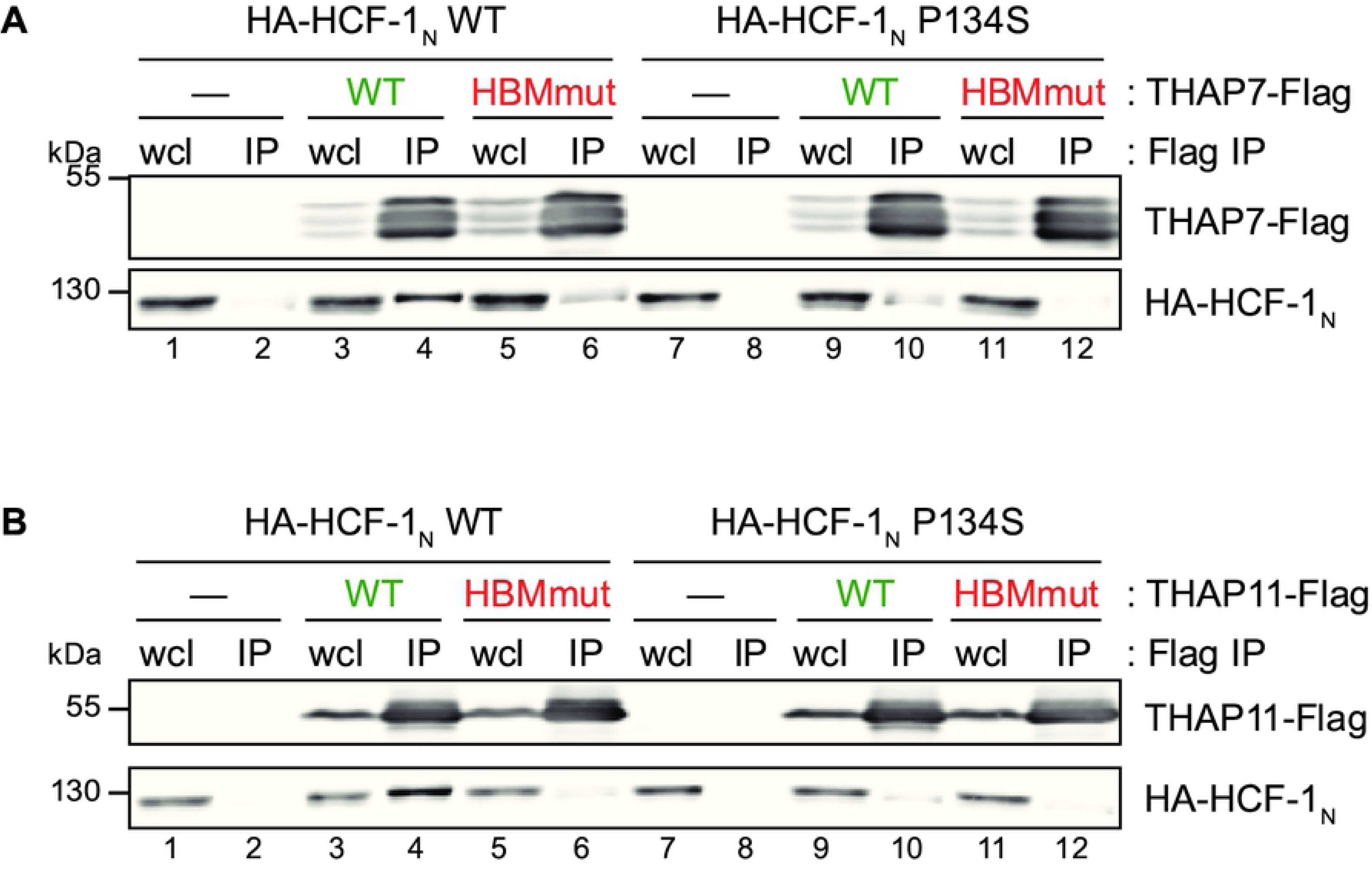
THAP7 and THAP11 interacts with HCF-1 via their respective HBM and Kelch domains. HEK-293 cells were co-transfected with WT or P134S forms of HA-HCF-1, with or without (A) THAP7-Flag or (B) THAP11-Flag constructs. Whole-cell lysates were subjected to Flag immunoprecipitation and analyzed by immunoblot with anti-Flag (upper panels) and anti-HA (lower panels) antibodies. HBM-positive THAP proteins are labeled in green whereas HBM-negative ones are in red. wcl, whole-cell lysate; IP, immunoprecipitate.

To characterize the HBM–Kelch domain contribution to these THAP-protein–HCF-1 interactions, we (i) took advantage of the well-characterized HCF-1 Kelch-domain point mutation called P134S, which prevents its association with HBM-containing target proteins [44], and (ii) created HBM-disruptive point mutants in THAP7 and THAP11. Each mutation on its own impairs the THAP-protein – HCF-1_N_ association (Figs 3A and B, compare lanes 4 with lanes 6 and 10) and when in combination further disrupt the interaction (lane 12). Consequently, these two THAP proteins both bind HCF-1 via the well-characterized interaction between the HCF-1 Kelch domain and HBM.

Thus, THAP7 and THAP11 form homodimers likely via their respective coiled-coil domains and associate with HCF-1 using their respective HBM sequences. Although these two interaction surfaces lie in proximity to one another in THAP proteins (e.g., by 9 amino acids in THAP11), at least one HBM in a double HA and Flag epitope-tagged THAP11 “heterodimer” can be disrupted without preventing THAP11 dimer formation as shown in Fig 2B (compare lanes 10 with 8); thus THAP-protein dimer formation may not be dependent on HCF-1 association.

### THAP7 promotes cell proliferation

To study the impact of THAP7 on cell proliferation, we engineered multiple homozygous clones of each of three point-mutant HEK-293 cell lines using CRISPR/Cas9 mediated genome editing [45,46] (S3 Fig and Materials and Methods): (i) a likely THAP7 null mutant (THAP7_null_), in which we created nonsense mutations at both codons 13 and 14 of the *THAP7* coding sequence to disrupt THAP7 synthesis (three clones, S3A Fig); (ii) the aforementioned THAP7 HBM mutant (THAP7_HBM_) (two clones, S3B Fig), and (iii) a THAP7 mutant C-terminally truncated after the HBM via two sequential nonsense mutations, thus deleting the coiled-coil domain (THAP7_ΔCC_) (four clones, S3C Fig).

The three THAP7_null_ mutant cell lines were viable albeit cell numbers increased more slowly than for the parental cells (labeled THAP7_WT_ in Fig 4). Visually, we did not observe an increase in dead cells and thus conclude that THAP7 is important for HEK-293-cell proliferation (i.e., cell-cycle progression). The two THAP7_HBM_ mutant clones proliferated at different rates (S4A Fig): one as the parental cells and the other as the THAP7_null_ mutant cells preventing any conclusion to be drawn regarding the impact of the HBM mutation on THAP7 function. In contrast, the four THAP7_ΔCC_ mutant clones (S4B Fig) consistently exhibited a slowdown in cell proliferation similar to that of the THAP7_null_ cells (compare Fig 4 and S4B Fig), suggesting that the THAP7 coiled-coil domain, and thus likely its ability to form dimers, is critical for the THAP7 function observed in HEK-293 cell proliferation.

**Fig 4.**
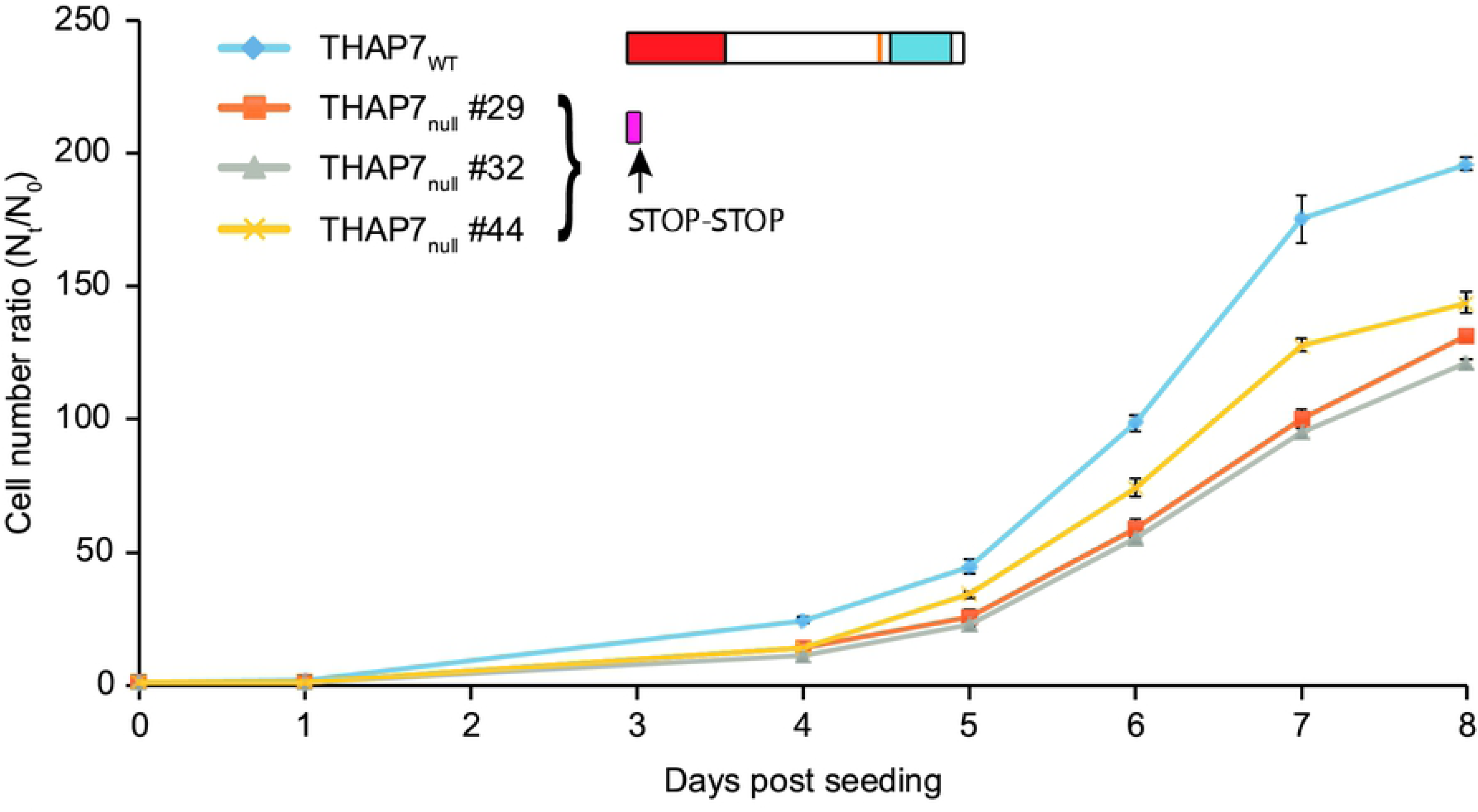
Disruption of the *THAP7* gene in HEK-293 cells impairs cell proliferation. THAP7_WT_ and three independent THAP7_null_ cell lines were seeded at the same density on day 0 (1.25 x 10^4^ cells per ml). For each cell line, 2 plates were used for counting every 24 hours from day 1 to day 8 (except days 2 and 3). The ratio of the mean cell counts between duplicates (N_t_) and the initial cell number (N_0_), with standard deviation, is plotted. Cartoons of the THAP7_WT_ and THAP7_null_ protein structures are shown.

### Translation of the THAP11_F80L_ cobalamin-disorder-associated mutation into a cellular model

We attempted to obtain THAP11 mutant cells corresponding to those described for THAP7, but no THAP11_null_ or THAP11_ΔCC_ cell clone was obtained. Furthermore, only one heterozygous THAP11_HBM_ mutant clone was obtained (S5A Fig) and these cells failed to survive. These results suggest that THAP11 is necessary for HEK-293-cell viability and that both the HBM and coiled-coil domain are important for THAP11 function.

During the course of this study, a human THAP11 mutation was identified in a patient with cobalamin disorder [22]. We thus took advantage of this finding to focus, as we do in the remainder of this study, on the clarification of the functional consequences of this cobalamin-disorder-associated THAP11_F80L_ mutation. The THAP11 F80 phenylalanine residue is highly conserved across vertebrate species [22] and also among the 12 different human THAP proteins (S1 Fig). As labeled in blue in S1 Fig, it is at the last position of the so-called “AVPTI<u>F</u>” box (green box in S1 Fig) lying at the C terminus of the THAP-domain; this box has been suggested to promote proper folding of the DNA-binding zinc finger [3,4]. We managed to obtain a single clone bearing the homozygous THAP11_F80L_-encoding C240G mutation (S5B Fig). Although still proliferating, the mutant cells increased in cell number at less than one-half the rate of the parental cells (Fig 5). As for the THAP7 mutant cells, we did not observe any increase in cell death, suggesting that THAP11 is important in sustaining HEK-293-cell proliferation.

**Fig 5.**
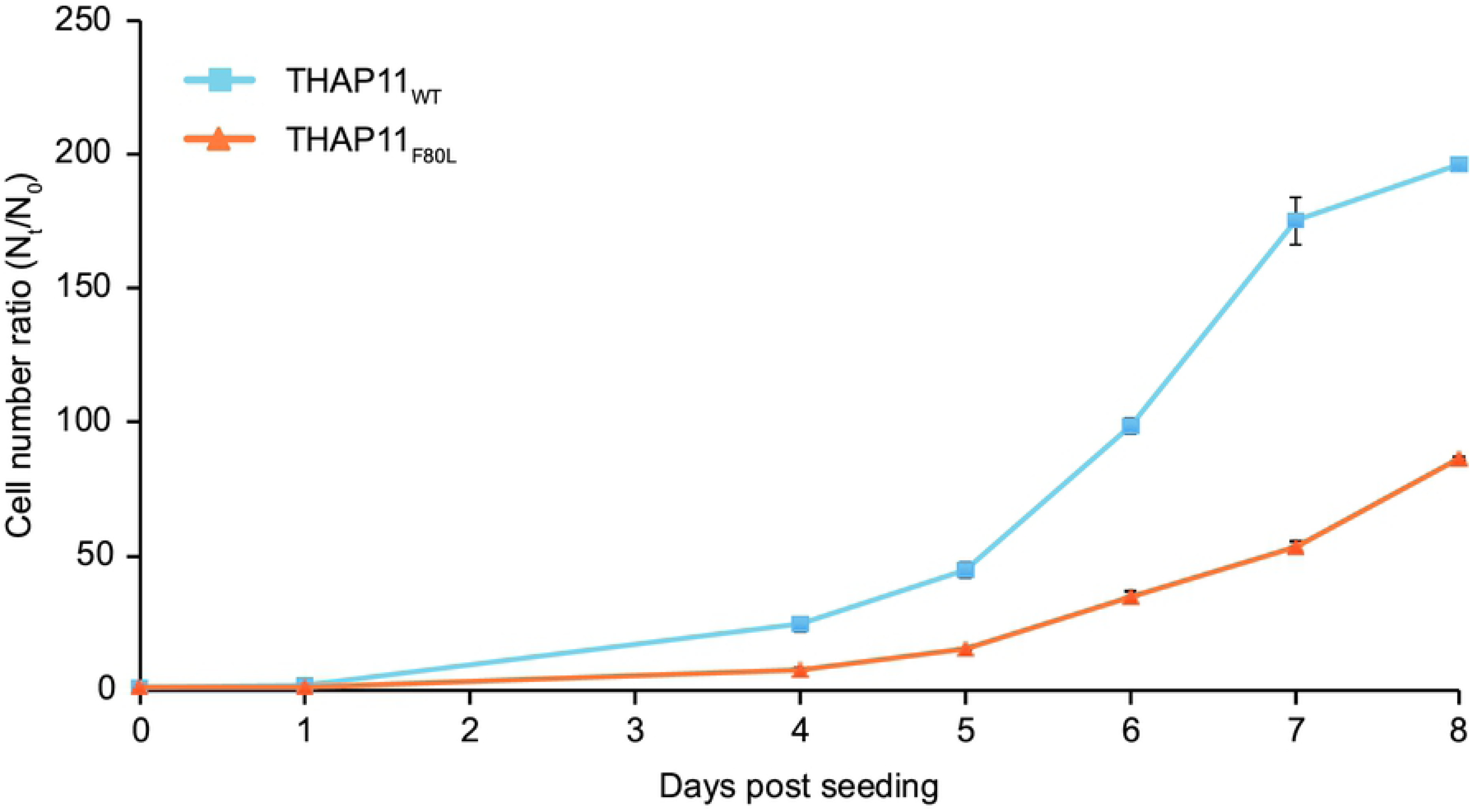
The THAP11_F80L_ mutation impairs HEK-293-cell proliferation. THAP11_WT_ and THAP11_F80L_ cells were seeded at the same density on day 0 (1.25 x 10^4^ cells per ml). For each cell line, 2 plates were used for counting every 24 hours from day 1 to day 8 (except days 2 and 3). The ratio of the mean cell counts between duplicates (N_t_) and the initial cell number (N_0_), with standard deviation, is plotted.

To determine the defect(s) caused by the THAP11_F80L_ mutation, we performed both THAP11-chromatin immunoprecipitation followed by high-throughput sequencing (ChIP-seq) and high-throughput RNA sequencing (RNA-seq) with both parental and THAP11_F80L_ mutant cells. Using this information, we first analyzed the effects of the THAP11_F80L_ mutation on THAP11 itself. As previously described [5], THAP11 binds to its own promoter at two near-by sites (overlapping THAP11-associated motifs; see below) of which only one was affected by the THAP11_F80L_ mutation (Fig 6A, track 2 and 4). Despite this difference in *THAP11*-promoter binding, consistent with what has been observed in fibroblasts from the *THAP11_F80L_* patient [22], *THAP11_F80L_* mRNA levels were unchanged relative to the *THAP11_WT_* levels (Fig 6B). In contrast, the levels of THAP11_F80L_ protein were reduced relative to the THAP11_WT_ protein (Fig 6C). The difference in signal is unlikely owing to an altered recognition of the mutant protein by the antibody, because the antibody used recognizes the last 85 amino acids of THAP11 distal to the F80L mutation. This difference in protein level could be explained by protein destabilization as the mutation affects the “AVPTIF” box implicated in THAP-domain stability. Whichever the case, the levels of THAP_F80L_ protein in the mutant cells does not obviously affect *THAP11* gene transcription.

**Fig 6.**
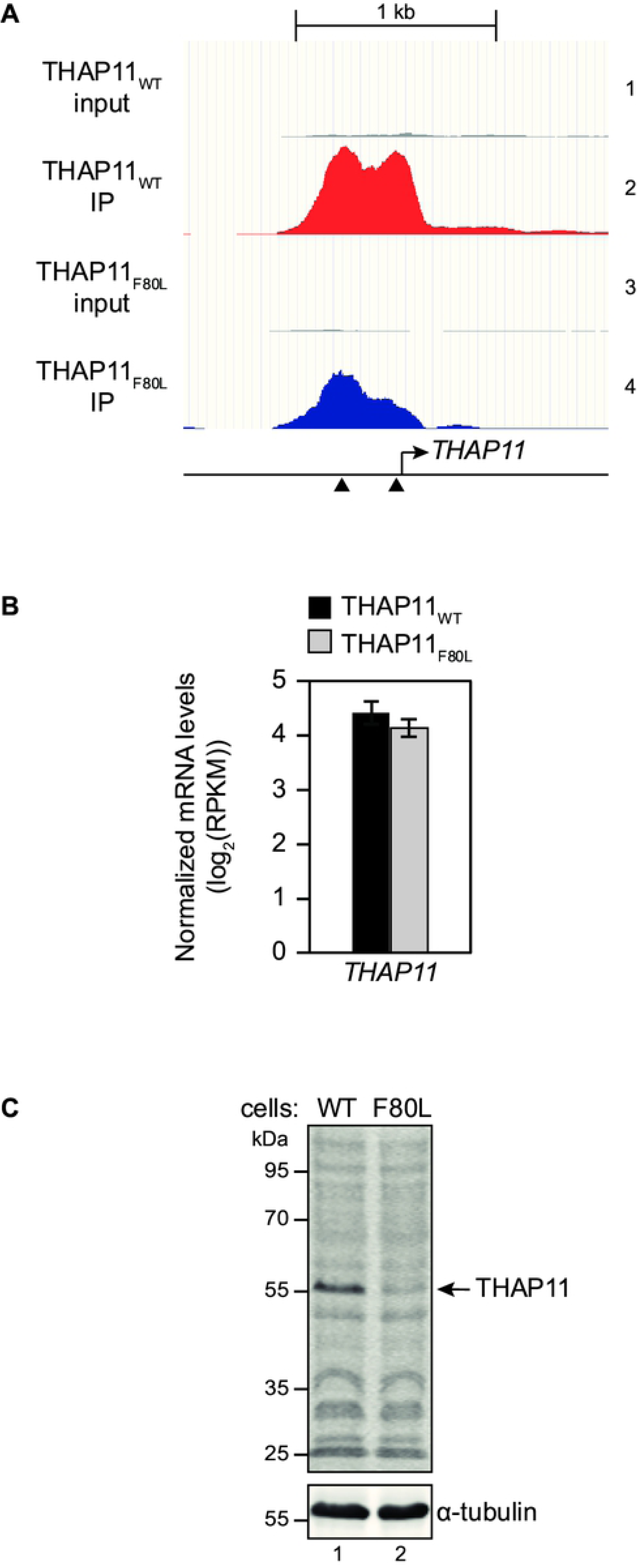
Effects of the THAP11_F80L_ mutation on *THAP11* expression. (A) THAP11 occupancy on its cognate gene promoter (chr16:67,875,200-67,877,300) in THAP11_WT_ (tracks 1 and 2) and THAP11_F80L_ cells (tracks 3 and 4). THAP11_WT_ (red) and THAP11_F80L_ (blue) ChIP fragment densities are visualized in parallel with their respective input (gray) — meaning the pre-IP total chromatin — densities (tracks 1 and 3, respectively), using the UCSC genome browser. All tracks are set with the same vertical viewing range. The two black triangles represent two THAP11-associated motifs (see Fig 7). (B) Normalized *THAP11* mRNA levels in THAP11_WT_ (black) and THAP11_F80L_ (gray) cells, shown as the mean +/- standard deviation of the *THAP11* log_2_(RPKM) of two biological replicates. (C) Equal protein amounts of THAP11_WT_ (lane 1) and THAP11_F80L_ (lane 2) whole cell lysates were analyzed by immunoblot using an anti-THAP11 antibody (top panel) and anti-α-tubulin as a loading control (bottom panel).

### The THAP11_F80L_ mutation selectively affects THAP11 DNA association

A global view (see Material and Methods for analysis parameters) of the THAP11 ChIP-seq results in parental and THAP11_F80L_ HEK-293 cells revealed 2341 THAP11_WT_ and only 1473 THAP11_F80L_ peaks, with about half of the peaks in each case being located within +/– 250 bp of an annotated transcription start site (TSS) (Fig 7A and S1 Table). S2 Table shows that the 1114 TSS-associated THAP11_WT_ peaks belong to expressed genes most prominently involved in transcription and its regulation, indicating the THAP11 controls programs of transcription, consistent with what has been previously reported [5].

**Fig 7.**
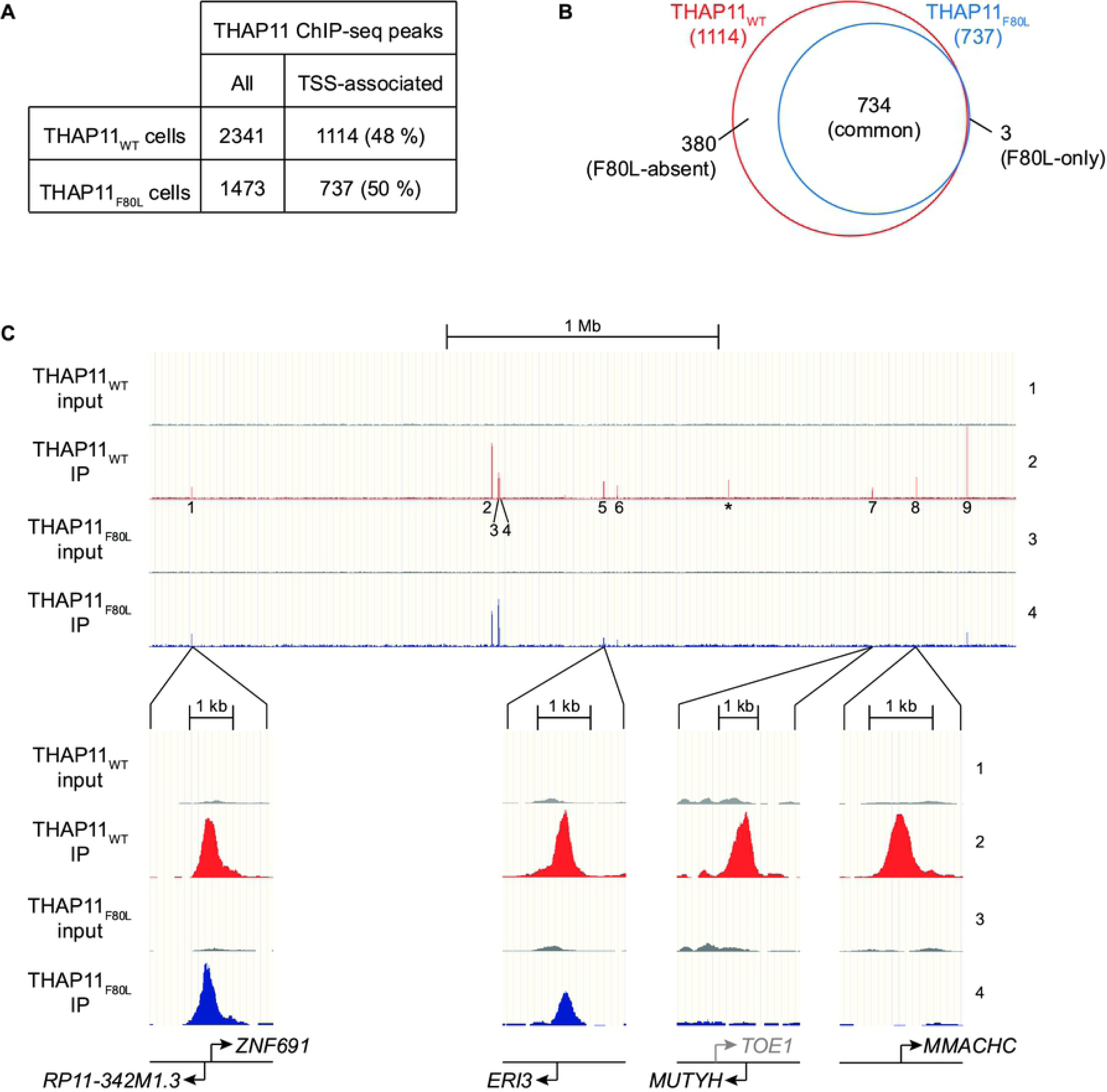
The THAP11_F80L_ mutation affects selectively THAP11 DNA association. (A) Number of peaks in THAP11_WT_ and THAP11_F80L_ cells: total peak number (left) and number and percentage of total peaks located within +/– 250 bp of an annotated TSS (right). (B) Venn diagram showing the overlap of THAP11_WT_ (red) and THAP11_F80L_ (blue) TSS-associated peaks. (C) THAP11 occupancy in a 3.3 Mb region of Chromosome 1 (chr1:43,100,000-46,400,000) including the *MMACHC* gene in THAP11_WT_ (tracks 1 and 2) and THAP11_F80L_ (tracks 3 and 4) cells. THAP11_WT_ (track 2, red) and THAP11_F80L_ (track 4, blue) ChIP fragment densities are visualized in parallel with their respective input densities (tracks 1 and 3, respectively; gray), using the UCSC genome browser. Each set of tracks is set with the same vertical viewing range — which differs from set to set — and the 9 TSS-associated THAP11_WT_ peaks are labelled (1-9). *, a non TSS-associated THAP11_WT_ peak. Close-ups of representative regions discussed in the text are shown below. Note in peak 7, the *TOE1* gene TSS is shown in gray because it falls outside of the +/– 250 bp THAP11-peak cut-off. Only protein-coding genes are displayed.

Being particularly interested in THAP11 regulation of transcription and to focus the study, we present the analysis of the TSS-associated THAP11 peaks. To characterize these peaks, we classified them into three categories (Fig 7B): (i) peaks present in both the THAP11_WT_ parental and the THAP11_F80L_ mutant samples called “common”, (ii) peaks missing (i.e., falling below the peak-calling threshold) in the THAP11_F80L_ sample called “F80L-absent”, and (iii) peaks present exclusively in the THAP11_F80L_ sample called “F80L-only”. As shown in the Venn diagram in Fig 7B, only three TSS-associated peaks fell into the “F80L-only” category (these peaks had a low significance score and were attributed to noise in the data; they were subsequently ignored), whereas 380 TSS-associated peaks were in the F80L-absent category — fully one-third of the total 1114 THAP11_WT_ peaks. Thus, the THAP11_F80L_ mutant protein binds to only a subset of the THAP11_WT_ promoter targets and does no exhibit any evident de novo promoter binding compared to the THAP11_WT_ protein.

Fig 7C illustrates THAP11_WT_ vs. THAP11_F80L_ DNA-site specificity with a 3.3 Mb view of Chromosome 1 that includes the *MMACHC* gene. The view covers nine THAP11_WT_ TSS-associated peaks (1–9) — of which four (peaks 1, 5, 7 and 8) were selected for close-ups to illustrate the points illuminated below — and one non-TSS-associated peak (labeled *): two TSS-associated peaks (peaks 7 and 8) and the one non-TSS-associated THAP11 peak fell into the THAP11_F80L_-absent category and the remainder in the common category (see S3 Table). Among those in the common category, we note that some peaks (e.g., peak 1 in Fig. 7C) remain largely the same size between the THAP11_WT_ and THAP11_F80L_ samples, whereas others (e.g., peak 5) were smaller albeit still present in the THAP11_F80L_ sample. In the subsequent analyses described below, we do not distinguish between these two common peak subcategories. In conclusion, the THAP11_F80L_ mutation results in a selective disruption of THAP11 DNA binding at specific promoters in HEK-293 cells, and does not create de-novo THAP11 promoter-binding sites.

### Restricted DNA-sequence recognition by the THAP11_F80L_ protein

To examine more broadly the nature of the effect of the THAP11_F80L_ mutation, we calculated size scores for each peak (see Materials and Methods for peak score determination) and plotted separately the distribution of the common and F80L-absent peak scores for both the THAP11_WT_ parental and THAP11_F80L_ mutant samples (Fig 8A). Consistent with their *a priori* categorization, the THAP11_F80L_ peaks scored higher in the common group than in the F80L-absent group (dashed lines). This tendency was also reflected at the level of the THAP11_WT_ scores as the common peaks (dark-red solid line) scored higher than the F80L-absent peaks (light-red solid line). The peak-score difference between the THAP11_WT_ and THAP11_F80L_ common peaks (dark-red lines) was less pronounced, however, than the difference between the THAP11_WT_ and THAP11_F80L_ F80L-absent peaks (light-red lines). Together, these observations suggest that: (i) the F80L-absent peaks represent lower affinity THAP11_WT_ binding sites and (ii) binding to these sites is hypersensitive to the THAP11_F80L_ mutation.

**Fig 8.**
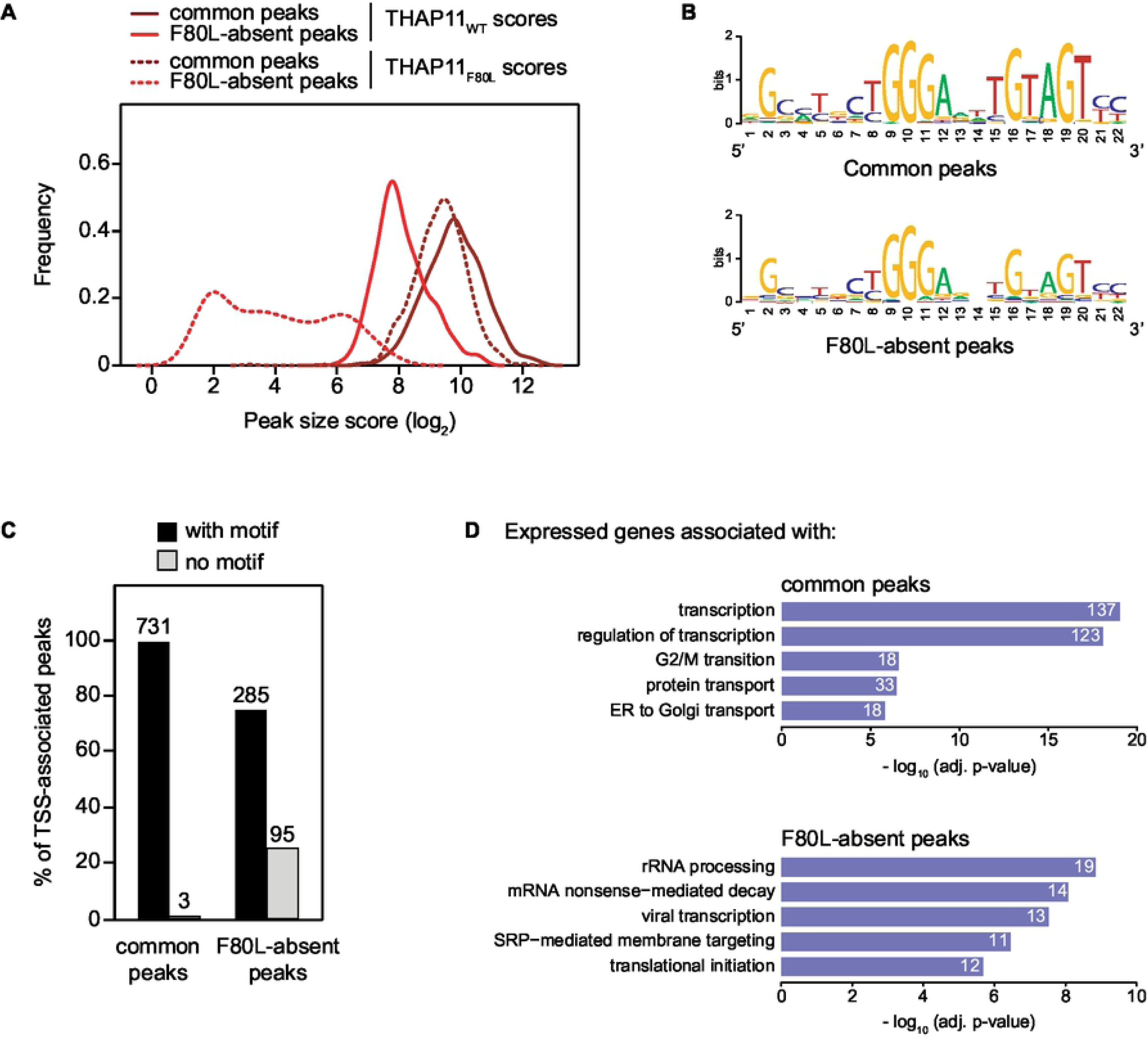
Restricted DNA-sequence recognition by the THAP11_F80L_ protein. (A) Distribution of the common (dark-red lines) and F80L-absent (light-red lines) peak scores for both the THAP11_WT_ parental (solid lines) and THAP11_F80L_ mutant (dashed lines) samples. Size scores were calculated for each peak (see Extended Materials and Methods in S1 File) and their frequency was plotted. (B) Motif logos of THAP11-associated motif (TAM) consensus sequences for the single closest TAM within +/– 250 bp to either the common (upper) or the F80L-absent (lower) peak summits. (C) Plot of the percentage of the common (left) and F80L-absent (right) peaks with one or more TAMs (black bars), or without a TAM (gray bars). Raw numbers are indicated above the bars. (D) Summary of the GO analyses of the expressed genes associated with the common (upper panel) and F80L-absent (lower panel) peaks; see S3 Table for the complete GO analysis results. Only the top-five GO biological process terms, with an adjusted p-value below 0.001, are shown. The number of expressed genes associated to each GO term is written on the corresponding bar.

To understand the reason for these differences, we examined the DNA sequences underlying the common and F80L-absent peaks. The most prominent motif to emerge from this analysis of both sets of peaks was the THAP11-associated motif (TAM) also referred to as RBM [5] or SBS2 [47]. We conclude that the DNA sequences under both common and F80L-absent peaks are enriched for the TAM, and thus that the difference between the two sets of peaks may result from subtle differences among TAMs.

We probed this possibility by separately generating TAM consensus sequences (Fig 8B) for the closest TAM associated with either the common or the F80L-absent peaks. Of the 22 bp-selected TAM sequence shown in Fig 8B, the common-peak-associated one (top) displays two prominent areas of consensus: a 5’ CTGGGA sequence and a 3’ TGTAGT sequence. The 5’ CTGGGA consensus was as prominent among the TAMs of the F80L-absent peaks, but the 3’ TGTAGT consensus was significantly weaker (Fig 8B, bottom). Taken together with the peak score analysis in Fig 8A, this comparison suggests that the THAP11_F80L_ protein is more sensitive to changes in the 3’ half of the consensus than the THAP11_WT_ protein.

In addition to a weaker TAM consensus sequence associated with the F80L-absent peaks, fully one-quarter of the F80L-absent peaks had no discernible nearby TAM (i.e., within 1000 bp on each side of the peak maximum), whereas essentially all common peaks were associated with one or more TAMs (Fig 8C). These analyses further emphasize the importance of a strong TAM consensus sequence for THAP11_F80L_ promoter recognition.

In addition to examining the TAMs of THAP11-peak-containing promoters, we asked if the nature of the genes associated with the common and F80L-absent peaks differ through gene-ontology (GO) analysis as summarized in Fig 8D (see S4 Table). Indeed, whereas the common peaks were prominently associated with transcription-regulatory genes, the association makeup of the F80L-absent-peak group was more heterogeneous and no longer evidently associated with transcription. Thus, a core function of THAP11 — to regulate programs of transcription — is apparently sufficiently robust to resist the effects of the THAP11_F80L_ mutation; it may be disruption of such a core function of THAP11 that makes the aforementioned THAP11_null_, THAP11_HBM_ and THAP11_ΔCC_ mutations lethal.

### Altered patterns of gene expression in THAP11_F80L_ cells

To examine the consequences of the THAP11_F80L_ mutation at the gene-expression level we analyzed the RNA-seq results (S5 Table). Figure 9A plots the transcript levels for protein-encoding genes in THAP11_WT_ cells (X axis) against their fold change in THAP11_F80L_ cells (Y axis). More often transcript levels considered to be changing (see fold change and p-value cutoffs in Materials and Methods) had decreased in THAP11_F80L_ cells compared to THAP11_WT_ cells: 523 genes were downregulated (red) whereas 279 genes were upregulated (blue). Furthermore, the magnitudes of decrease were greater than those for increases. Thus, the THAP11_F80L_ mutation apparently affects activation of transcription — either directly or indirectly — more often than repression.

**Fig 9.**
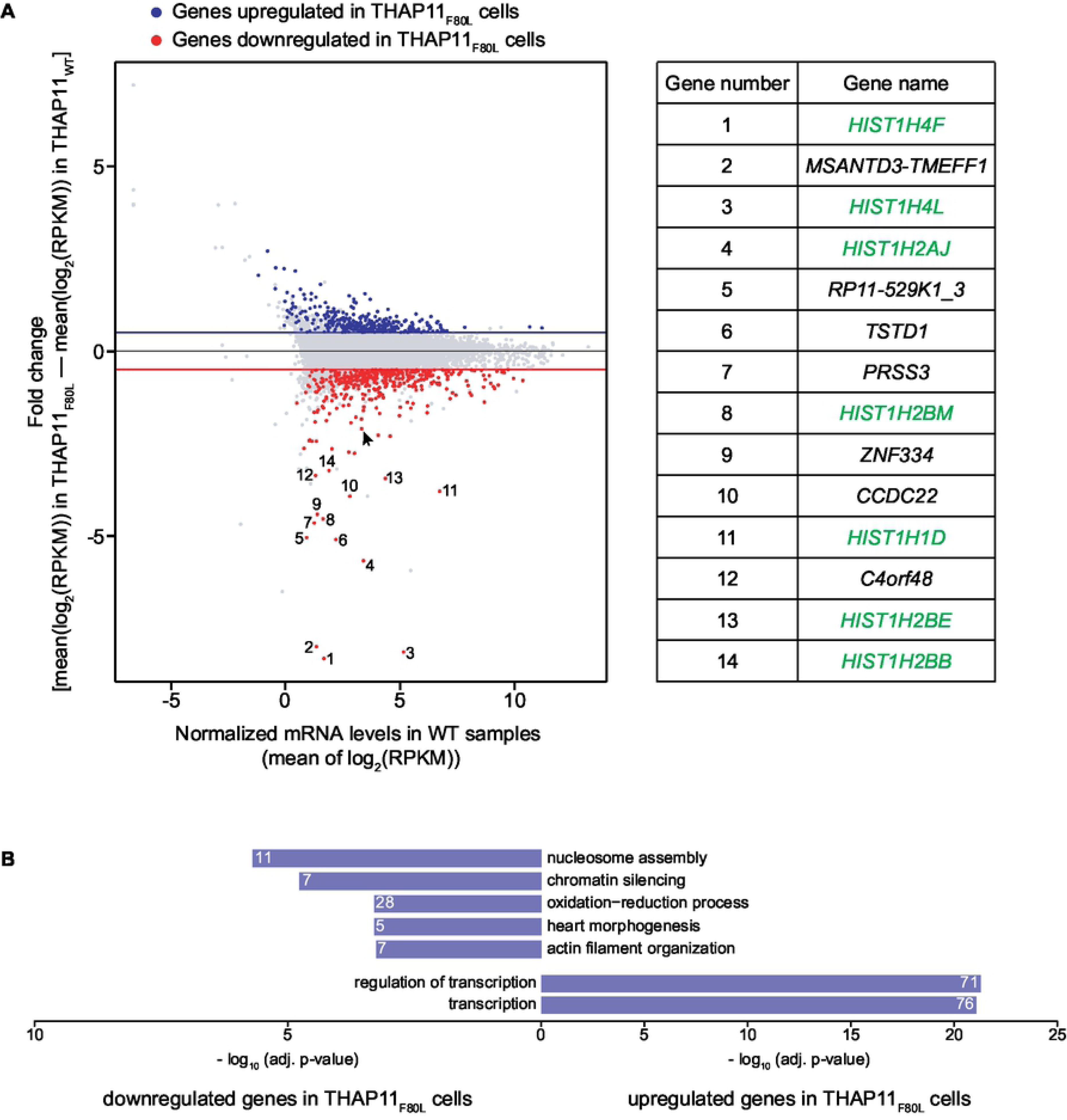
Altered patterns of gene expression in THAP11_F80L_ cells. (A) The transcript levels for protein-encoding genes in THAP11_WT_ cells (X axis) was plotted against their fold change in THAP11_F80L_ cells (Y axis). Red, downregulated genes; blue, upregulated genes; gray, genes with an adjusted p-value above 0.05 or an absolute fold-change below 0.5. Arrow, *MMACHC* gene. The 14 genes most downregulated and with an adjusted p-value below 0.05 are numbered and listed (right) with canonical S-phase histone-encoding genes in green. (B) Summary of the separate GO analyses of the upregulated (lower) and downregulated (upper) protein-coding genes — see S6 Table for the complete GO analysis results. Only the top-five GO biological process terms having an adjusted p-value below 0.001 are shown. The number of changing genes associated to each GO term is written on the corresponding bar.

Among the 14 genes most downregulated in the THAP11_F80L_ cells, fully one-half were canonical S-phase histone-encoding genes (Fig. 9A, right). In our THAP11_WT_ ChIP-seq data, the THAP11 protein was absent from canonical histone-encoding gene promoters (S1 Table) indicating that this downregulation is an indirect effect. Such downregulation may be linked to the slower proliferation rate of THAP11_F80L_ cells, either as a cause or a consequence.

Separate GO analyses of the upregulated and downregulated genes (Fig 9B and S6 Table) revealed enrichment for different gene functions. The downregulated set was more heterogeneous with weaker enrichments for each GO term than the upregulated set. Consistent with the aforementioned histone-gene expression change, it included nucleosome assembly and chromatin silencing terms. In contrast, the upregulated set was associated with only two GO terms that fell below the p-value cutoff — these being both transcription oriented and with highly significant p-values. Thus, whereas more genes were downregulated in THAP11_F80L_ cells, those that were upregulated were strongly enriched for a single biological process, transcription. Although the THAP11_F80L_ ChIP-seq results indicated that THAP11 association with the TSS of transcription-related genes was overall resistant to the F80L mutation, the THAP11_F80L_ mutation appears to nevertheless affect THAP11 regulation of gene-transcription programs.

### Likely direct roles of THAP11 affected by the THAP11_F80L_ mutation

To probe which of these gene-transcription changes might be direct consequences of THAP11_F80L_ function, we identified those changing genes whose promoters were directly bound by THAP11_WT_ but not THAP11_F80L_ (i.e., F80L-absent ChIP-seq peaks). These genes included 11 upregulated and 44 downregulated genes as shown in the Fig 10 heat map (see also S3 Table for *MMACHC* and *MUTYH*). By GO analysis (S7 Table), we observed that a significant portion (one quarter) of these direct-effect genes are associated with regulation of transcription (Fig 10, blue arrows). This observation suggests that THAP11 plays a higher-order role in the regulation of gene transcription by directly regulating — most often activating — the expression of secondary transcriptional regulator-encoding genes.

**Fig 10.**
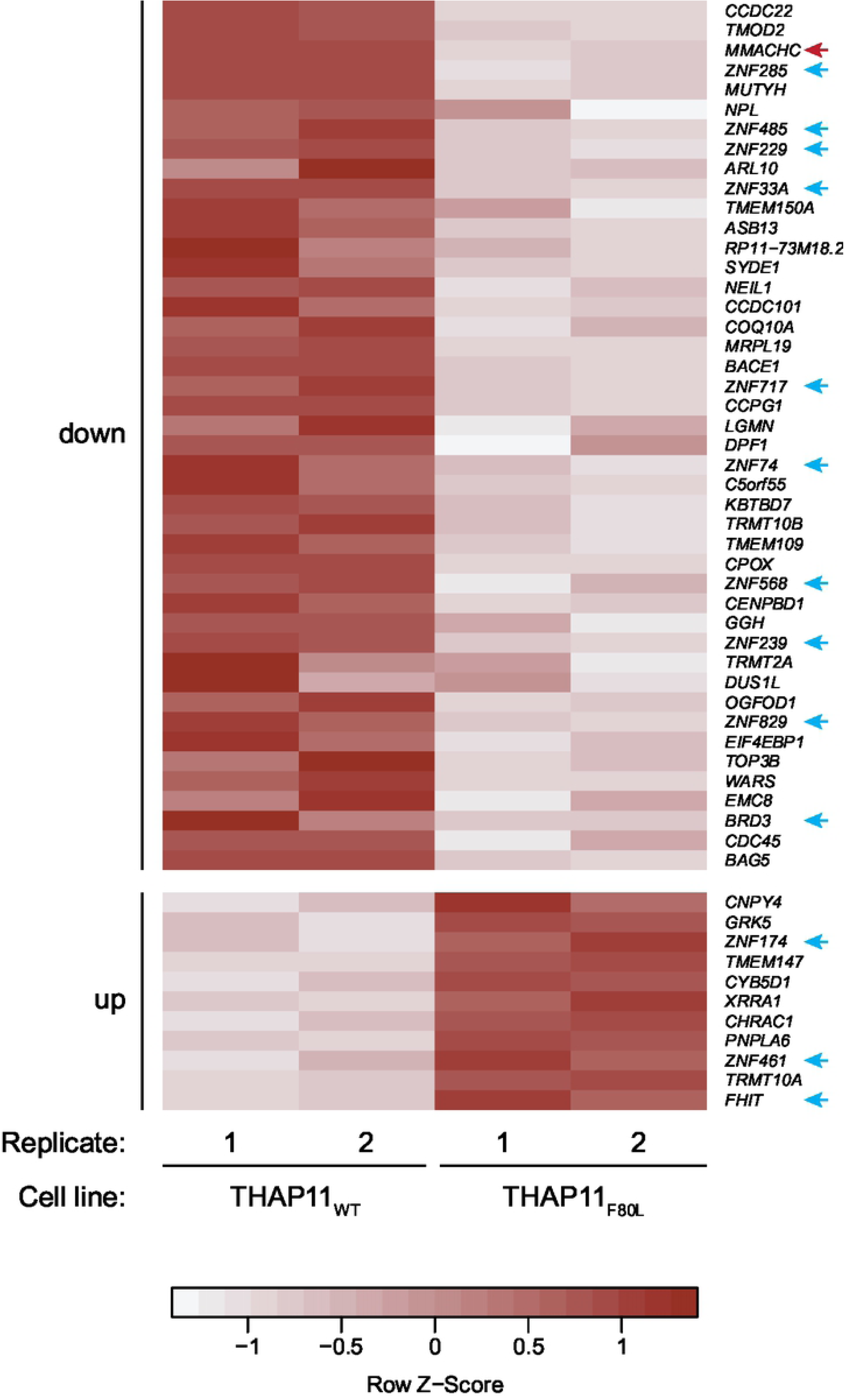
Direct gene expression changes caused by the THAP11_F80L_ mutation. Gene-expression value heat map of the 11 upregulated (bottom) and 44 downregulated (upper) protein-coding genes whose promoters were directly bound by THAP11_WT_ but not THAP11_F80L_ (i.e., F80L-absent ChIP-seq peaks). The gene-expression scores for replicates 1 and 2 are indicated for both the THAP11_WT_ (left) and the THAP11_F80L_ (right) cells and the genes are ranked by fold change, from the highest to the lowest, separately for the downregulated (top) from the upregulated (bottom) genes. Blue arrows, genes associated with the regulation of transcription GO term (see S7 Table); red arrow, *MMACHC* gene.

Fig 10 also shows that the *MMACHC* gene ranks third among the most downregulated direct-target genes in THAP11_F80L_ cells (red arrow). This observation suggests that THAP11 activates *MMACHC* gene transcription and that the THAP11_F80L_ mutation impairs this activation, consistent with the cobalamin-disorder phenotype of the patient from whom the mutation was identified [22].

## Discussion

We have studied the THAP family of DNA-binding proteins extending from an evolutionary perspective of the 12 human members to a detailed analysis of a human disease associated THAP11 mutation.

### The THAP family of proteins

The sequence comparison in Fig 1 accentuates how the THAP family is rapidly evolving — even just between human and mouse there are five additional *THAP* genes in human. Nevertheless, two non-THAP domain sequence elements, the coiled-coil domain and HBM, are shared among the large majority of the 12 human THAP proteins, illuminating their importance for this class of proteins.

The coiled-coil domain is implicated in association of THAP proteins [11,48,49]. As described previously [11,41] and reproduced here, THAP11 forms homodimers. We show that THAP7 also forms homodimers. In contrast, we did not detect human THAP7 and THAP11 heterodimer formation, and thus did not observe a previously reported THAP7 association with an N-terminal THAP11 protein fragment [11]. A wider study (S8 Table and [40]) has shown homodimer formation for THAP8 and heterodimer formation between THAP8 and THAP11 but not heterodimer formation between THAP4 and THAP11 proteins. Thus, THAP homo- and heterodimer formation possibilities are likely complex, permitting complicated transcriptional regulatory networks. This complexity is probably amplified by the rapid evolution of divergent DNA recognition properties of individual THAP domains [3–7].

The HBM permits many THAP proteins to extend their regulatory network to association with the transcription co-regulator HCF-1. We have shown that both THAP7 and THAP11 interact with HCF-1 via their respective HBM. Nevertheless, the presence or absence of an HBM in a THAP protein does not determine whether it will or will not associate with HCF-1: THAP5 contains an HBM but failed to associate with HCF-1, and THAP8 lacks an HBM and yet associated with HCF-1 (S8 Table and [40]). These results further underscore the complex regulatory potential of the THAP-protein family.

### Roles of THAP7 and THAP11 in cell proliferation and viability

To probe regulatory roles of THAP proteins, in particular in cell proliferation, we designed targeted mutations for the *THAP7* and *THAP11* genes in HEK-293 cells to create null mutations and HBM and coiled-coil deficient proteins. For *THAP7*, we obtained all three categories of mutation, in contrast to *THAP11* where we only transiently obtained a heterozygous HBM mutant. The sensitivity of HEK-293 cells to *THAP11* null mutagenesis is consistent with the embryonic lethality observed upon mouse *Thap11* (Ronin) deletion [11]. Indeed, THAP11 is even sensitive to individual HBM and coiled-coil disruption, indicating critical roles of THAP11 interaction with HCF-1 and dimer formation for cell viability.

Although THAP7 was not essential for HEK-293-cell viability, we observed consistent retarded cell proliferation for the THAP7_null_ and THAP7_ΔCC_ mutant cells. Although we did not probe for apoptosis directly, no increase in dying cells was observed during culturing, suggesting that the decrease in cell accumulation reflects a defect in cell-cycle progression. Thus, these two THAP proteins appear to play important and in one case — THAP11 — an essential role in HEK-293-cell viability.

### The THAP11_F80L_ cobalamin-disorder associated mutation

We observed that *THAP11* is particularly sensitive to mutagenesis, as we did not obtain any viable THAP11_null_, THAP11_HBM_ or THAP11_ΔCC_ mutants. This mutational sensitivity is also highlighted by the ExAC database of human exon sequence variants (http://exac.broadinstitute.org/gene/ENSG00000168286, [50]) where (i) the only six loss-of-function mutations reported are all located in the *THAP11* poly-glutamine tract-encoding sequences and (ii) no missense mutations were reported in the *THAP11* sequences encoding the THAP domain. This restricted number of genetic variants altering the THAP11 protein highlights the importance of THAP11 functions. And yet an individual homozygous for the THAP11_F80L_ mutation survived for 10 years [22], showing how a *THAP11* mutation, located in the THAP domain, can still be tolerated for viability.

We took advantage of this finding to study the function of a natural human THAP11 mutant. We observed decreased cell proliferation in HEK-293 THAP11_F80L_ mutant cells. We also noted that, although *THAP11_F80L_* and *THAP11_WT_* mRNA levels were similar, there was less THAP11_F80L_ protein in the mutant cells compared to THAP11_WT_ protein in unmutated cells. The THAP11_F80L_ mutation affects a residue of the “AVPTIF” box, which is suggested to contribute to proper folding of the zinc finger [3,4]. The location of the mutation and the lower levels of THAP11_F80L_ protein — without concomitant decrease in *THAP11_F80L_* mRNA — suggest that the mutant THAP11_F80L_ protein is destabilized and more rapidly degraded.

To investigate the consequences of the THAP11_F80L_ mutation, we probed THAP11 DNA binding using THAP11 ChIP-seq. We observed no significant new THAP11 TSS-associated-binding sites, but did observe that the mutation causes the loss of THAP11 DNA binding at a specific subset of TSS-associated sites. A detailed analysis of the altered binding-site patterns revealed that some TSS-associated sites are particularly sensitive to the THAP11_F80L_ mutation. Overall, these are sites exhibiting a lower affinity for THAP11 binding, with a weaker THAP11 motif consensus sequence — particularly at the 3’ half of the consensus sequence. These observations suggest that THAP11-binding sites come in two classes that may respond to wild-type THAP11 activity differently, for example by being more or less sensitive to the activity of regulatory co-factors or dimer formation.

The analysis of how the THAP11_F80L_ mutation affects gene expression revealed alterations in the regulation of programs of transcription. The gene-expression changes can be direct or indirect consequences of the THAP11_F80L_ mutation and subsequent changes in DNA binding. For instance, the impaired expression of numerous canonical S-phase histone-encoding genes in THAP11_F80L_ cells is likely the result of an indirect effect of the mutation, as THAP11 is absent from the corresponding histone promoters. Indeed, the decrease of histone-gene expression could be the result, not the cause, of the decrease in cell proliferation observed in these mutant cells. Alternatively, the THAP11_F80L_ mutation and the alteration of THAP11 DNA-binding activity could have direct consequences for the expression of specific genes, apparently especially for genes implicated in the regulation of transcription.

### Linking the THAP11_F80L_ mutation with human disease

Consistent with identification in a patient with cobalamin disorder [22], the THAP11_F80L_ mutation also has an apparent direct effect on the activity of the *MMACHC* promoter. Whereas THAP11_WT_ binds to the *MMACHC* promoter, the THAP11_F80L_ mutant binds poorly to the *MMACHC* promoter, consistent with the reduction in *MMACHC* mRNA levels observed in the fibroblasts of the patient in which the mutation was isolated [22]. These results suggest a direct link, missing so far, between the THAP11_F80L_ mutation and the cobalamin disorder observed in the associated patient.

Interestingly, the corresponding F-to-L missense mutation has also been identified in human THAP1, also named DYT6. This THAP1_F81L_ mutation is one of the THAP1 mutations associated with dystonia 6 [51]. Dystonia 6, a subtype of primary monogenic torsion dystonia, is a movement disorder involving dysfunctions of the central nervous system and characterized by involuntary muscle contractions. It is caused by a variety of THAP1 mutations (90 to date) spread over the entire *THAP1* gene [23–25,52,53]. Studies are conflicting as to the effect of the THAP1_F81L_ mutation on THAP1 DNA binding — with either little, if any [54], partial [51], or apparently complete [55] loss of THAP1-DNA binding described, suggesting that the effect of the mutation on DNA binding is sensitive to the conditions for analysis. For example, as for THAP11_F80L_, the THAP1_F81L_ protein is destabilized [54], which could influence DNA-binding affinity *in vivo*. Thus, the matching mutations in two distinct THAP proteins with different gene targets and cellular roles [3–7], have similar effects on the protein — consistent with the identical disruption of a highly conserved protein segment — and yet very different phenotypic and clinical effects.

The different results presented here suggest a general model for how the THAP11_F80L_ mutation is pathogenic: the mutation affects THAP11 protein levels and perhaps, by virtue of this position in the DNA-binding THAP domain, its DNA-binding activities. Either of these attributes could lead to the observed bimodal specificity of promoter occupancy by THAP11_F80L_ protein. We imagine that often the less-abundant mutated THAP11_F80L_ protein retains binding activity for DNA sites of strongest affinity — e.g., the ones with the strongest TAM consensus sequence.

This general model, however, is unlikely to explain all the effects of the THAP11_F80L_ mutation, because some promoters, in particular the *MMACHC* promoter, can be greatly affected by the THAP11_F80L_ mutation but possess TAMs with good matches to the TAM consensus sequence. And yet it is precisely the great sensitivity of the *MMACHC* promoter to the THAP11_F80L_ mutation that likely accounts in large part for the associated cobalamin-disorder phenotype, with other gene expression changes likely accounting for additional clinical manifestations that provide the specificity of this cobalamin-disorder subtype [33].

Importantly, the results described here indicate a direct role of the THAP11_F80L_ mutation in *MMACHC* gene expression and thus in the cobalamin disorder diagnosed in the patient.

## Materials and methods

Detailed materials and methods are described in Supporting Information S1 File.

### Cell proliferation assays

Cells were seeded at the same density (1.25 x 10^4^ cells per ml) on day 0, and for each cell line, 2 plates used for counting every 24 hours from day 1 to day 8 (except days 2 and 3). THAP7_HBM_, THAP7_ΔCC_, and parental cell lines were tested in parallel in a first experiment, whereas THAP11_F80L_, THAP7_null_ and parental cell lines were tested in parallel in a second experiment.

### Immunoblotting

The samples were separated by SDS-PAGE before being transferred onto a nitrocellulose membrane. After blocking, membranes were incubated overnight at 4°C with the primary antibody. After washing, they were further incubated with the appropriate secondary antibody during 1 hour at room temperature. Blots were finally visualized with the Odyssey R infra-red imaging system (LI-COR).

### CRISPR/Cas9 mutagenesis

CRISPR/Cas9-mediated mutagenesis in HEK 293 cells was performed as recommended [56] with minor modifications. After co-transfection of the cells with a GFP, Cas9 and gRNA encoding plasmid together with repair template, cells were cell sorted and single-cell clones were tested by restriction digest. The mutation was further confirmed by sequencing.

### RNA extraction and sequencing (RNA-seq)

Total RNA was used to prepare ribosomal RNA-depleted libraries for 50-nucleotide single-read high-throughput sequencing using an Illumina HiSeq 2100 device with 6 samples per lane (multiplexing).

### Chromatin immunoprecipitation and sequencing (ChIP-seq)

For each ChIP, 5 IPs were performed in parallel and subsequently pooled, each time by incubating 9 μg of crosslinked and sonicated chromatin at 4°C with 2 μg of anti-THAP11 antibody and further with protein G agarose beads. Pre-IP samples (total chromatin — input sample) were prepared in parallel. 5 ng of each ChIP and input purified DNA were used to prepare paired-end sequencing libraries for 100-nucleotide paired-end high-throughput sequencing (Illumina, HiSeq 2100) with 3 samples per lane (multiplexing).

### Bioinformatics

#### Analysis of THAP proteins

The evolutionary tree (Fig 1) was prepared by multiple sequence alignment (Muscle alignment) on the 90 N-terminal amino-acids of each human THAP protein sequence. The alignment was refined and visualized using the iTOL (interactive Tree Of Life) online tool [57,58].

#### RNA-seq analysis

Single reads were mapped onto the Hg19 human genome annotation using STAR (Spliced Transcripts Alignment to a Reference, [59]) and read counts and normalized RPKM (Reads Per Kilobase of transcript per Million mapped reads) were calculated using RSEM [60,61]. Only protein-coding genes were considered and genes with an RPKM value below 1.2 in all the samples were considered as not expressed and discarded from further analyses. Differential analyses were performed with DESeq2 [62]. Resulting gene sets were used for Gene Ontology (GO) enrichment analysis [63,64].

#### ChIP-seq analysis

Paired-end reads were mapped onto the Hg19 human genome annotation using STAR (Spliced Transcripts Alignment to a Reference) [59]. Peaks were detected using the Model-based Analysis of ChIP-Seq (MACS2) tool [65] and tested using the “Origami method” (V. Praz; see [66]), which identified enriched bins. Only MACS-identified peaks intersecting with Origami-enriched bins were kept. A peak was considered TSS associated if one or more underlying nucleotides was located within +/- 250 bp of an RNA polymerase II transcription start site. Peak scores were defined as the difference of the IP and input log_2_ counts, scaled by total tags for the sample and the peak width. Peaks were visualized with the UCSC genome browser [67].

#### Data availability

The datasets used in this study are available in the NCBI Gene Expression Omnibus (GEO, http://www.ncbi.nlm.nih.gov/geo) under the Super Series accession number GSE138208.

## Acknowledgments

We thank the Lausanne Genomic Technologies Facilities for preparation of the RNA-seq libraries and the high-throughput sequencing; the EPFL Flow Cytometry Core Facility for cell sorting; Laura Sposito for advice on HEK-293-cell CRISPR/Cas9 targeted mutagenesis; Fabienne Lammers for technical advice; Liliane Michalik and Jolanda van Leeuwen for critical readings of the manuscript.

## Supporting information

**S1 File. Extended experimental methods.** This file describes in detail the materials and methods used in the experiments reported in this article.

**S1 Fig. The THAP domain of human THAP proteins.** Sequence alignment of the first 90 amino-acids of the 12 human THAP proteins, with the conserved features highlighted as follows: red, “C2CH signature” of the zinc finger; purple, strictly conserved residues; green box, “AVPTIF box”; blue, phenylalanine residue mutated in the THAP11_F80L_ cobalamin-disorder patient.

**S2 Fig. THAP7 is phosphorylated and endogenous THAP11 interacts with HCF-1 via the HCF-1 N-terminal subunit.** (A) HEK-293 cells were transfected with THAP7-Flag construct, and whole-cell lysate (lane 1) was either directly treated with calf intestinal phosphatase (CIP) (lane 2) or subjected to Flag immunoprecipitation (lane 3) before being treated with CIP (lanes 4 and 5), and analyzed by immunoblot with an anti-Flag antibody. d.CIP, heat-inactivated CIP. Relative to Fig 2B. (B) HEK-293 cells were transfected without (lanes 1 and 2) or with HA-HCF-1_C_ (lanes 3 and 4), HA-HCF-1_N_ (lanes 5 and 6), or HA-HCF-1_FL_ (lanes 7 and 8) constructs and whole-cell lysates (lanes 1, 3, 5, and 7) subjected to HA immunoprecipitation (lanes 2, 4, 6, and 8) and analyzed by immunoblot with anti-HA (two upper panels) and anti-THAP11 (lower panel) antibodies. Relative to Fig 3B. wcl, whole-cell lysate; IP, immunoprecipitate.

**S3 Fig. THAP7 CRISPR/Cas9 mutants.** Details of the mutagenesis (left) and sequencing chromatograms (right) of the (A) THAP7_null_, (B) THAP7_HBM_, and (C) THAP7_ΔCC_ mutant clones. The mutated nucleotides and resulting amino-acids are depicted in red in the mutant sequences.

**S4 Fig. Effect of the THAP7_HBM_ and THAP7_ΔCC_ mutations on HEK-293-cell proliferation.** THAP7_WT_ and (A) two independent THAP7_HBM_ or (B) four independent THAP7_ΔCC_ cell lines were seeded at the same density (1.25 x 10^4^ cells per ml) on day 0, and for each cell line, 2 plates used for counting every 24 hours from day 1 to day 8 (except days 2 and 3). The ratio of the mean cell counts between duplicates (N_t_) and the initial cell number (N_0_), with standard deviation, is plotted. Cartoons of the THAP7_WT_, THAP7_HBM_ and THAP7_ΔCC_ protein structures are shown. Relative to Fig 4.

**S5 Fig. THAP11 CRISPR/Cas9 mutants.** Details of the mutagenesis (left) and sequencing chromatograms (right) of the (A) THAP11_HBM_ and (B) THAP11_F80L_ mutant clones. The mutated nucleotides and resulting amino-acids are depicted in red in the mutant sequences.

**S1 Table. List of ChIP-seq peaks.** Table listing the peaks identified in the ChIP-seq experiment (all peaks, and not only TSS-associated peaks). Each peak has been identified with a unique identifier (column A) and categorized as “common”, “F80L absent” or “F80L only” (see text. Column B). The exact peak position is detailed in columns D and E (genomic coordinates of the start and the end of the peak, respectively). The peak scores and counts in the THAP11_WT_ (columns F and H) and THAP11_WT_ (columns G and I) peaks are indicated. Details about the THAP11-associated motifs are indicated: total number of motifs in a region expanding 1000 bp on each side of the peak maximum (column J), genomic coordinates of the start (column K) and end (column L) of the closest motif to the peak center, motif sequence (column M), motif E-value relative to the consensus motif (column N) and the relative position of the motif to the peak (column O). Details of the genes identified under the peaks are listed, together with their RNA-seq data: number of genes having their TSS in a region expanding 250 bp on each side of the peak boundaries (column P), distance of the TSS gene to the peak (columns R, AB, AL and AV), gene strand (columns S, AC, AM and AW), gene type (columns T, AD, AN and AX), normalized gene mRNA levels (log_2_(RPKM)) in each of the THAP11_WT_ (columns U and V; AE and AF; AO and AP; AY and AZ) and THAP11_F80L_ (columns W and X; AG and AH; AQ and AR; BA and BB) biological replicates, the (log_2_) THAP11_F80L_ versus THAP11_WT_ fold change and associated adjusted p-value of gene expression values (columns Y and Z; AI and AJ; AS and AT; BC and BD). NA, non-applicable, meaning no such feature (motif of gene) relative to the peak. ND, non-determined, meaning gene not expressed in our dataset.

**S2 Table. THAP11 controls programs of transcription.** Complete results of the KEGG, GO, BIOCARTA and REACTOME analyses performed on the 1138 expressed genes associated with the 1114 TSS-associated THAP11_WT_ peaks.

**S3 Table. List of THAP11 peaks around the *MMACHC* gene promoter.** Features of the 10 THAP11 peaks in a 3.3 Mb region of Chromosome 1 including the *MMACHC* gene. Relative to Fig 7C.

**S4 Table. Nature of the genes associated with the common and F80L-absent peaks.** Complete results of the KEGG, GO, BIOCARTA and REACTOME analyses performed on the expressed genes associated with the common (first sheet) or F80L-absent (second sheet) peaks. Relative to Fig 8D.

**S5 Table. RNA-seq data analysis of THAP11_WT_ and THAP11_F80L_ HEK-293 cells.** Table listing the expressed genes identified in the RNA-seq experiment, with their respective expression values. Each gene is identified by its Ensembl ID (column A) and its name (column B), the gene type and description being detailed (columns C and D, respectively). The results of the RNA-seq data are presented for all expressed genes (see Materials and Methods for details) as follows: normalized counts of each biological replicate in THAP11_WT_ (columns E and F) and THAP11_F80L_ (columns G and H) cells; log_2_(RPKM) values of each biological replicate in THAP11_WT_ (columns I and J) and THAP11_F80L_ (columns K and L) cells; (log_2_) THAP11_F80L_ versus THAP11_WT_ fold change (column M); and associated p-values (column N) and adjusted p-value (column O).

**S6 Table. Nature of the genes differently expressed between the THAP11_WT_ and THAP11_F80L_ cells.** Complete results of the KEGG, GO, BIOCARTA and REACTOME analyses performed on the expressed genes downregulated (first sheet) or upregulated (second sheet) in the THAP11_F80L_ cells. Relative to Fig 9B.

**S7 Table. Direct gene expression changes caused by the THAP11_F80L_ mutation.** Complete results of the KEGG, GO, BIOCARTA and REACTOME analyses performed on the expressed genes changing in the mutant THAP11_F80L_ cells whose promoter is directly bound by THAP11_WT_ but not THAP11_F80L_. Relative to Fig 10.

**S8 Table. Summary of the results of the wider study investigating THAP dimerization and interaction with HCF-1.** Green plus, interaction; red minus, no interaction; (—), very weak, if any, interaction; grey dot, not tested. See [40].

## References

1. Roussigne M, Kossida S, Lavigne A-C, Clouaire T, Ecochard V, Glories A, et al. The THAP domain: a novel protein motif with similarity to the DNA-binding domain of P element transposase. Trends Biochem Sci. 2003;28: 66–9. doi:10.1016/S0968-0004(02)00013-0

2. Clouaire T, Roussigne M, Ecochard V, Mathe C, Amalric F, Girard J-P. The THAP domain of THAP1 is a large C2CH module with zinc-dependent sequence-specific DNA-binding activity. Proc Natl Acad Sci U S A. 2005;102: 6907–12. doi:10.1073/pnas.0406882102

3. Bessière D, Lacroix C, Campagne S, Ecochard V, Guillet V, Mourey L, et al. Structure-function analysis of the THAP zinc finger of THAP1, a large C2CH DNA-binding module linked to Rb/E2F pathways. J Biol Chem. 2008;283: 4352–4363. doi:10.1074/jbc.M707537200

4. Gervais V, Campagne S, Durand J, Muller I, Milon A. NMR studies of a new family of DNA binding proteins: The THAP proteins. J Biomol NMR. 2013;56: 3–15. doi:10.1007/s10858-012-9699-1

5. Dejosez M, Levine SS, Frampton GM, Whyte WA, Stratton SA, Barton MC, et al. Ronin/Hcf-1 binds to a hyperconserved enhancer element and regulates genes involved in the growth of embryonic stem cells. Genes Dev. 2010;24: 1479–1484. doi:10.1101/gad.1935210

6. Sabogal A, Lyubimov AY, Corn JE, Berger JM, Rio DC. THAP proteins target specific DNA sites through bipartite recognition of adjacent major and minor grooves. Nat Struct Mol Biol. 2010;17: 117–124. doi:10.1038/nsmb.1742

7. Balakrishnan MP, Cilenti L, Ambivero C, Goto Y, Takata M, Turkson J, et al. THAP5 is a DNA-binding transcriptional repressor that is regulated in melanoma cells during DNA damage-induced cell death. Biochem Biophys Res Commun. 2011;404: 195–200. doi:10.1016/j.bbrc.2010.11.092

8. Macfarlan T, Parker JB, Nagata K, Chakravarti D. Thanatos-associated protein 7 associates with template activating factor-Ibeta and inhibits histone acetylation to repress transcription. Mol Endocrinol. 2006;20: 335–47. doi:10.1210/me.2005-0248

9. Macfarlan T, Kutney S, Altman B, Montross R, Yu J, Chakravarti D. Human THAP7 is a chromatin-associated, histone tail-binding protein that represses transcription via recruitment of HDAC3 and nuclear hormone receptor corepressor. J Biol Chem. 2005;280: 7346–7358. doi:10.1074/jbc.M411675200

10. Miele A, Medina R, Van Wijnen AJ, Stein GS, Stein JL. The interactome of the histone gene regulatory factor HiNF-P suggests novel cell cycle related roles in transcriptional control and RNA processing. J Cell Biochem. 2007;102: 136–148. doi:10.1002/jcb.21284

11. Dejosez M, Krumenacker JS, Zitur LJ, Passeri M, Chu LF, Songyang Z, et al. Ronin Is Essential for Embryogenesis and the Pluripotency of Mouse Embryonic Stem Cells. Cell. 2008;133: 1162–1174. doi:10.1016/j.cell.2008.05.047

12. Parker JB, Palchaudhuri S, Yin H, Wei J, Chakravarti D. A Transcriptional Regulatory Role of the THAP11-HCF-1 Complex in Colon Cancer Cell Function. Mol Cell Biol. 2012;32: 1654–1670. doi:10.1128/MCB.06033-11

13. Zhu CY, Li CY, Li Y, Zhan YQ, Li YH, Xu CW, et al. Cell growth suppression by thanatos-associated protein 11(THAP11) is mediated by transcriptional downregulation of c-Myc. Cell Death Differ. 2009;16: 395–405. doi:10.1038/cdd.2008.160

14. Nakamura S, Yokota D, Tan L, Nagata Y, Takemura T, Hirano I, et al. Down-regulation of Thanatos-associated protein 11 by BCR-ABL promotes CML cell proliferation through c-Myc expression. Int J Cancer. 2012;130: 1046–1059. doi:10.1002/ijc.26065

15. Parker JB, Yin H, Vinckevicius A, Chakravarti D. Host cell factor-1 recruitment to E2F-bound and cell-cycle-control genes is mediated by THAP11 and ZNF143. Cell Rep. 2014;9: 967–82. doi:10.1016/j.celrep.2014.09.051

16. Michaud J, Praz V, Faresse NJ, JnBaptiste CK, Tyagi S, Schütz F, et al. HCFC1 is a common component of active human CpG-island promoters and coincides with ZNF143, THAP11, YY1, and GABP transcription factor occupancy. Genome Res. 2013;23: 907–916. doi:10.1101/gr.150078.112

17. Fujita J, Freire P, Coarfa C, Benham AL, Gunaratne P, Schneider MD, et al. Ronin Governs Early Heart Development by Controlling Core Gene Expression Programs. Cell Rep. 2017;21: 1562–1573. doi:10.1016/j.celrep.2017.10.036

18. Durruthy-Durruthy J, Wossidlo M, Pai S, Takahashi Y, Kang G, Omberg L, et al. Spatiotemporal Reconstruction of the Human Blastocyst by Single-Cell Gene-Expression Analysis Informs Induction of Naive Pluripotency. Dev Cell. 2016;38: 100– 115. doi:10.1016/j.devcel.2016.06.014

19. Seifert BA, Dejosez M, Zwaka TP. Ronin influences the DNA damage response in pluripotent stem cells. Stem Cell Res. 2017;23: 98–104. doi:10.1016/j.scr.2017.06.014

20. Kong XZ, Yin RH, Ning HM, Zheng WW, Dong XM, Yang Y, et al. Effects of THAP11 on erythroid differentiation and megakaryocytic differentiation of K562 cells. PLoS One. 2014;9. doi:10.1371/journal.pone.0091557

21. Poché RA, Zhang M, Rueda EM, Tong X, McElwee ML, Wong L, et al. RONIN Is an Essential Transcriptional Regulator of Genes Required for Mitochondrial Function in the Developing Retina. Cell Rep. 2016;14: 1684–1697. doi:10.1016/j.celrep.2016.01.039

22. Quintana AM, Yu HC, Brebner A, Pupavac M, Geiger EA, Watson A, et al. Mutations in THAP11 cause an inborn error of cobalamin metabolism and developmental abnormalities. Hum Mol Genet. 2017;26: 2838–2849. doi:10.1093/hmg/ddx157

23. Ozelius LJ, Bressman SB. Genetic and clinical features of primary torsion dystonia. Neurobiol Dis. 2011;42: 127–135. doi:10.1016/j.nbd.2010.12.012

24. Bragg DC, Armata IA, Nery FC, Breakefield XO, Sharma N. Molecular pathways in dystonia. Neurobiol Dis. 2011;42: 136–147. doi:10.1016/j.nbd.2010.11.015

25. LeDoux MS, Xiao J, Rudzińska M, Bastian RW, Wszolek ZK, Van Gerpen JA, et al. Genotype-phenotype correlations in THAP1 dystonia: Molecular foundations and description of new cases. Park Relat Disord. 2012;18: 414–425. doi:10.1016/j.parkreldis.2012.02.001

26. Balakrishnan MP, Cilenti L, Mashak Z, Popat P, Alnemri ES, Zervos AS. THAP5 is a human cardiac-specific inhibitor of cell cycle that is cleaved by the proapoptotic Omi/HtrA2 protease during cell death. Am J Physiol Heart Circ Physiol. 2009;297: H643–53. doi:10.1152/ajpheart.00234.2009

27. Li Y, Ning Q, Shi J, Chen Y, Jiang M, Gao L, et al. A novel epigenetic AML1-ETO/THAP10/miR-383 mini-circuitry contributes to t(8;21) leukaemogenesis. EMBO Mol Med. 2017;9: 933–949. doi:10.15252/emmm.201607180

28. Gladitz J, Klink B, Seifert M. Network-based analysis of oligodendrogliomas predicts novel cancer gene candidates within the region of the 1p/19q co-deletion. Acta Neuropathol Commun. 2018;6: 49. doi:10.1186/s40478-018-0544-y

29. Abate F, da Silva-Almeida AC, Zairis S, Robles-Valero J, Couronne L, Khiabanian H, et al. Activating mutations and translocations in the guanine exchange factor VAV1 in peripheral T-cell lymphomas. Proc Natl Acad Sci. 2017;114: 764–769. doi:10.1073/pnas.1608839114

30. de Souza Santos E, de Bessa SA, Netto MM, Nagai MA. Silencing of LRRC49 and THAP10 genes by bidirectional promoter hypermethylation is a frequent event in breast cancer. Int J Oncol. 2008;33: 25–31.

31. Johnson RA, Wright KD, Poppleton H, Mohankumar KM, Finkelstein D, Pounds SB, et al. Cross-species genomics matches driver mutations and cell compartments to model ependymoma. Nature. 2010;466: 632–636. doi:10.1038/nature09173

32. Lian WX, Yin RH, Kong XZ, Zhang T, Huang XH, Zheng WW, et al. THAP11, a novel binding protein of PCBP1, negatively regulates CD44 alternative splicing and cell invasion in a human hepatoma cell line. FEBS Lett. 2012;586: 1431–1438. doi:10.1016/j.febslet.2012.04.016

33. Sloan JL, Carrillo N, Adams D, Venditti CP. Disorders of Intracellular Cobalamin Metabolism. GeneReviews®. 2018. Available: http://www.ncbi.nlm.nih.gov/pubmed/20301503

34. Freiman RN, Herr W. Viral mimicry: Common mode of association with HCF by VP16 and the cellular protein LZIP. Genes Dev. 1997;11: 3122–3127. doi:10.1101/gad.11.23.3122

35. Lu R, Yang P, Padmakumar S, Misra V. The herpesvirus transactivator VP16 mimics a human basic domain leucine zipper protein, luman, in its interaction with HCF. J Virol. 1998;72: 6291–6297.

36. Burkhard P, Stetefeld J, Strelkov S V. Coiled coils: A highly versatile protein folding motif. Trends Cell Biol. 2001;11: 82–88. doi:10.1016/S0962-8924(00)01898-5

37. Sanghavi HM, Mallajosyala SS, Majumdar S. Classification of the human THAP protein family identifies an evolutionarily conserved coiled coil region. 2019; 2–11.

38. Lupas A, Van Dyke M, Stock J. Predicting coiled coils from protein sequences. Science. 1991;252: 1162–4. doi:10.1126/science.252.5009.1162

39. McDonnell A V, Jiang T, Keating AE, Berger B. Paircoil2: improved prediction of coiled coils from sequence. Bioinformatics. 2006;22: 356–8. doi:10.1093/bioinformatics/bti797

40. Dehaene H. THAP proteins in the transcriptional control of cell proliferation. Doctoral dissertation. University of Lausanne. 2019. Available: https://serval.unil.ch/notice/serval:BIB_02ABC667C655

41. Cukier CD, Maveyraud L, Saurel O, Guillet V, Milon A, Gervais V. The C-terminal region of the transcriptional regulator THAP11 forms a parallel coiled-coil domain involved in protein dimerization. J Struct Biol. 2016;194: 337–346. doi:10.1016/j.jsb.2016.03.010

42. Wilson AC, LaMarco K, Peterson MG, Herr W. The VP16 accessory protein HCF is a family of polypeptides processed from a large precursor protein. Cell. 1993;74: 115– 125. doi:10.1016/0092-8674(93)90299-6

43. Kristie TM, Pomerantz JL, Twomey TC, Parent SA, Sharp PA. The cellular C1 factor of the herpes simplex virus enhancer complex is a family of polypeptides. J Biol Chem. 1995;270: 4387–94. doi:10.1074/jbc.270.9.4387

44. Wilson AC, Freiman RN, Goto H, Nishimoto T, Herr W. VP16 targets an amino-terminal domain of HCF involved in cell cycle progression. Mol Cell Biol. 1997;17: 6139–46. Available: http://www.ncbi.nlm.nih.gov/pubmed/9315674

45. Jinek M, Chylinski K, Fonfara I, Hauer M, Doudna JA, Charpentier E. A programmable dual-RNA-guided DNA endonuclease in adaptive bacterial immunity. Science. 2012;337: 816–21. doi:10.1126/science.1225829

46. Costa Y, Ding J, Theunissen TW, Faiola F, Hore TA, Shliaha P V., et al. NANOG-dependent function of TET1 and TET2 in establishment of pluripotency. Nature. 2013;495: 370–374. doi:10.1038/nature11925

47. Ngondo-Mbongo RP, Myslinski E, Aster JC, Carbon P. Modulation of gene expression via overlapping binding sites exerted by ZNF143, Notch1 and THAP11. Nucleic Acids Res. 2013;41: 4000–4014. doi:10.1093/nar/gkt088

48. Sengel C, Gavarini S, Sharma N, Ozelius LJ, Bragg DC. Dimerization of the DYT6 dystonia protein, THAP1, requires residues within the coiled-coil domain. J Neurochem. 2011;118: 1087–1100. doi:10.1111/j.1471-4159.2011.07386.x

49. Richter A, Hollstein R, Hebert E, Vulinovic F, Eckhold J, Osmanovic A, et al. In-depth Characterization of the Homodimerization Domain of the Transcription Factor THAP1 and Dystonia-Causing Mutations Therein. J Mol Neurosci. 2017;62: 11–16. doi:10.1007/s12031-017-0904-2

50. Lek M, Karczewski KJ, Minikel E V., Samocha KE, Banks E, Fennell T, et al. Analysis of protein-coding genetic variation in 60,706 humans. Nature. 2016;536: 285–291. doi:10.1038/nature19057

51. Fuchs T, Gavarini S, Saunders-Pullman R, Raymond D, Ehrlich ME, Bressman SB, et al. Mutations in the THAP1 gene are responsible for DYT6 primary torsion dystonia. Nat Genet. 2009;41: 286–288. doi:10.1038/ng.304

52. Frederick NM, Shah P V, Didonna A, Langley MR, Kanthasamy AG, Opal P. Loss of the dystonia gene Thap1 leads to transcriptional deficits that converge on common pathogenic pathways in dystonic syndromes. Hum Mol Genet. 2019;28: 1343–1356. doi:10.1093/hmg/ddy433

53. Xiromerisiou G, Houlden H, Scarmeas N, Stamelou M, Kara E, Hardy J, et al. THAP1 Mutations And Dystonia Phenotypes : Genotype Phenotype Correlations. 2012;27: 1290–1294. doi:10.1002/mds.25146

54. Campagne S, Muller I, Milon A, Gervais V. Towards the classification of DYT6 dystonia mutants in the DNA-binding domain of THAP1. Nucleic Acids Res. 2012;40: 9927– 9940. doi:10.1093/nar/gks703

55. Gavarini S, Cayrol C, Fuchs T, Lyons N, Ehrlich ME, Girard JP, et al. Direct interaction between causative genes of DYT1 and DYT6 primary dystonia. Ann Neurol. 2010;68: 549–553. doi:10.1002/ana.22138

56. Ran FA, Hsu PD, Wright J, Agarwala V, Scott DA, Zhang F. Genome engineering using the CRISPR-Cas9 system. Nat Protoc. 2013;8: 2281–2308. doi:10.1038/nprot.2013.143

57. Letunic I, Bork P. Interactive Tree Of Life (iTOL): an online tool for phylogenetic tree display and annotation. Bioinformatics. 2007;23: 127–8. doi:10.1093/bioinformatics/btl529

58. Letunic I, Bork P. Interactive Tree Of Life v2: online annotation and display of phylogenetic trees made easy. Nucleic Acids Res. 2011;39: W475–8. doi:10.1093/nar/gkr201

59. Dobin A, Davis CA, Schlesinger F, Drenkow J, Zaleski C, Jha S, et al. STAR: ultrafast universal RNA-seq aligner. Bioinformatics. 2013;29: 15–21. doi:10.1093/bioinformatics/bts635

60. Li B, Dewey CN. RSEM: accurate transcript quantification from RNA-Seq data with or without a reference genome. BMC Bioinformatics. 2011;12: 323. doi:10.1186/1471-2105-12-323

61. Li B, Ruotti V, Stewart RM, Thomson JA, Dewey CN. RNA-Seq gene expression estimation with read mapping uncertainty. Bioinformatics. 2010;26: 493–500. doi:10.1093/bioinformatics/btp692

62. Love MI, Huber W, Anders S. Moderated estimation of fold change and dispersion for RNA-seq data with DESeq2. Genome Biol. 2014;15: 550. doi:10.1186/s13059-014-0550-8

63. The Gene Ontology Consortium. Expansion of the Gene Ontology knowledgebase and resources. Nucleic Acids Res. 2017;45: D331–D338. doi:10.1093/nar/gkw1108

64. Ashburner M, Ball CA, Blake JA, Botstein D, Butler H, Cherry JM, et al. Gene ontology: tool for the unification of biology. The Gene Ontology Consortium. Nat Genet. 2000;25: 25–9. doi:10.1038/75556

65. Zhang Y, Liu T, Meyer CA, Eeckhoute J, Johnson DS, Bernstein BE, et al. Model-based analysis of ChIP-Seq (MACS). Genome Biol. 2008;9: R137. doi:10.1186/gb-2008-9-9-r137

66. Renaud M, Praz V, Vieu E, Florens L, Washburn MP, L’Hôte P, et al. Gene duplication and neofunctionalization: POLR3G and POLR3GL. Genome Res. 2014;24: 37–51. doi:10.1101/gr.161570.113

67. Kent WJ, Sugnet CW, Furey TS, Roskin KM, Pringle TH, Zahler AM, et al. The Human Genome Browser at UCSC. Genome Res. 2002;12: 996–1006. doi:10.1101/gr.229102

